# Coexistence Maintained by Sexual Interactions

**DOI:** 10.64898/2026.01.27.702089

**Authors:** Adina F. Fébert, Géza Meszéna

## Abstract

Recent empirical studies have uncovered the coexistence of cryptic species that are morphologically indistinguishable yet genetically different. As they lack known ecological differentiation between them, it has been suggested that species-specific sexual interactions may be able to stabilize their coexistence. We investigated two types of such interactions: (1) traumatic mating suffered by females and (2) interference competition among males. We have learned that the consequences of the two types of sexual interactions are markedly different.

Traumatic mating may affect the fertility and/or survival of females. In both cases it introduces negative density dependence and, alone, can regulate the growth of a single population. Also, traumatic mating is able to stabilize the coexistence of ecologically equivalent populations, provided that it is stronger within species than between them. In contrast, male competition cannot constrain population growth without density dependence of a different origin. Accordingly, it destabilizes, rather than stabilizes, neutral coexistence of ecologically equivalent species.

A targeted, species-specific study of traumatic mating suffered by females would provide empirical support for the diversity-maintaining role of sexual interactions.

## Introduction

Cryptic species are widespread in wildlife. They have been reported in most types of organism and habitat, from fungi to freshwater fish, and from arthropods to arctic plants. Biologists have been aware of their existence for approximately 300 years, but only the recently developed DNA sequencing was able to solve the challenge of their taxonomical classification (Bickford et al. 2007). Lacking known niche differentiation, they represent a challenge for coexistence theory.

The traditional, reference interpretation of species coexistence is that it is stabilized by the negative frequency dependence caused by ecological niche segregation (Hutchinson 1957, 1959, 1978; MacArthur and Levins 1967). In this vein, one can assume that cryptic species coexist via ‘cryptic’ niche differentiation that is not reflected in morphology and is waiting for future identification. Alternatively, possibility of neutral coexistence among ecologically equivalent species can be considered. Such coexistence is necessarily temporal and expected to disappear through drift (McPeek, 2022, p. 344–363).

In this paper we are interested in a third possibility proposed first by Zhang et al. (1998) and several authors since: coexistence stabilized by sexual interactions. Gómez-Llano et al. (2021) reviewed these suggestions and classified sexual interactions based on whether they occur within or between sexes, or within or between species. These distinctions frame the present work also.

While some of the proposed mechanisms involve spatial structuring of the populations (see e.g. Kohda, 1995, 1998; Mikami et al., 2004), we study the simplest situations with spatial homogeneity. Such coexistence requires negative frequency dependence induced by sexual interactions (see e.g. Yamamichi et al., 2023). Specifically, we study two promising candidates mechanisms:

- Sexual conflict between sexes: *female harm* caused by traumatic mating, when the excessive or aggressive mating attempts of males affect females’ fertility, survival or both.
- Within sex interaction: Male-male interference competition, *male competition* for short. Both of these interactions are present in a wide range of species, and have been detected in relation to cryptic species. Female harm appears e.g., in case of fruit flies (*Drosophila*) and bed bugs (*Cimex lectularius*). *Drosophila* males tend to poison females slightly in order to prevent them from getting fertilized by another male (Rice et al., 2006), and *Cimex lectularius* females often suffer harm from males during copulation which is proved to decrease their fitness (Reinhardt et al., 2003). This latter mechanism also appears in case of spider species, (e.g. *Harpactea sadistica*, Rezác, 2009).

Male competition is displayed for example in case of cichlid fish (Kohda, 1995, 1998) and damselflyes, especially in the *Calopteryx* genus (Svensson et al., 2018). The males of these species are proven to repulse conspecifics harder than heterospecifics, which comes into view in context of their coexistence and, in the case of the latter, their richness of sympatric species – though spacial gradients and old genetic variation also tend to have an effect on this (Seehausen, 2015).

We investigate the emergence of frequency dependence from sexual interactions mechanistically, with explicit representation of the two sexes and their interactions. This approach is new, as total population sizes were the dynamical variables in the earlier models and the effects of sexual interactions on these variables were postulated on intuitive basis. We will see that the mechanistic underpinning leads to a deeper and more reliable understanding.

We proceed in two steps. First we ask, whether female harm, or male competition is able to regulate a single population. Then we investigate, whether these sexual interactions can stabilize the otherwise neutral coexistence of two ecologically equivalent species. We conclude with an outlook on the theoretical possibilities for cryptic coexistence and the possible ways to distinguish between them empirically.

Our simulation results presented in the Results section are supported analytically in the Appendix.

## Model definition

### Single species

At first, we study the dynamical consequences of the sexual interactions in a single sexually reproducing population. Females depend on males for reproduction and produce female and male offspring with equal ratio. On the other hand, overabundance of males may decrease fertility, or increase mortality of females. Male-male interference competition may increase male mortality, as well.

We describe the dynamics of the population with the following system of differential equations:

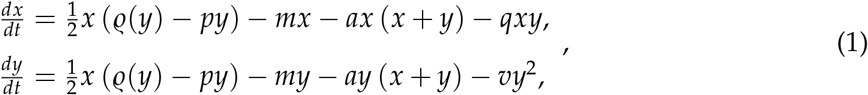

where *x* and *y* denotes abundance of females and males, respectively. Male dependence of female fertility rate is described by the saturating fertility function *ϱ*(*y*). Density-independent mortality is denoted by *m*. Strength of ecological regulation (e.g. exploitative competition) is represented by *a*. These variables are the same for both sexes. Coefficients *p* and *q* parametrize the two components of female harm caused by males: decreased fertility and increased mortality, respectively. Coefficient *v* denotes the intensity of male-male interference competition. Necessarily, *m* > 0, but the other coefficients can be 0 at some model versions.

We make the following assumptions about the fertility function:

- *ϱ*(*y*) is an increasing function of *y* ≥ 0 with *ϱ*(0) = 0 and *ϱ*_max_ = lim_*y*→∞_ *ϱ*(*y*) *<* ∞.
- Derivative *ϱ*′(*y*) > 0 is decreasing, that is, *ϱ*^*′′*^(*y*) *<* 0.

We will refer to *ϱ*′(*y*), as *Allee sensitivity*. The assumptions guaranty that

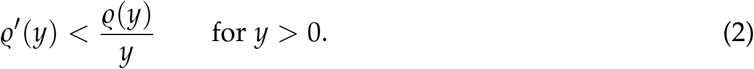

and

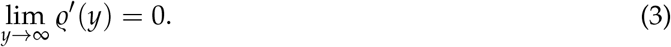

For the numerical investigations the fertility function *ϱ*(*y*) is specified, as a Holling type II functional response:

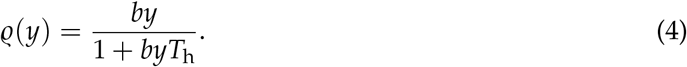

Here, *b* is the rate of search, e.g. the size of the territory searched by the searching party looking for a non-moving mating partner in unit time. The “handling time” *T*_h_ is the time needed after mating for a female to be ready to mate again. (We assume no handling time for males.)

Holling type II functional response can also be written in the usual form of the MichaelisMenten kinetics

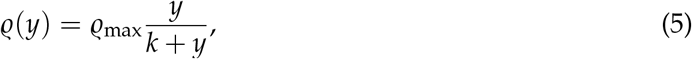

where 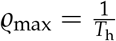 and 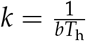 is the half-saturation constant.

The ODEs were integrated by Python’s built-in integrator *scipy*.*integrate*.*odeint*. The larger part of the analytic investigations in the Supplementary Material relies only on the qualitative saturating nature of the fertility function, as specified above.

### Two species

We investigate the joint dynamics of two non-hybridizing species, A and B, as specified above. They have a common ecological regulation and interact sexually. Both female harm and male competition can operate within, as well as between the species. That is, we assume that interspecific mating attempts may occur, therefore females may suffer force-copulation from allospecific males. Nevertheless, these matings do not produce viable offsprings.

The dynamics is specified by the following system of ODEs:

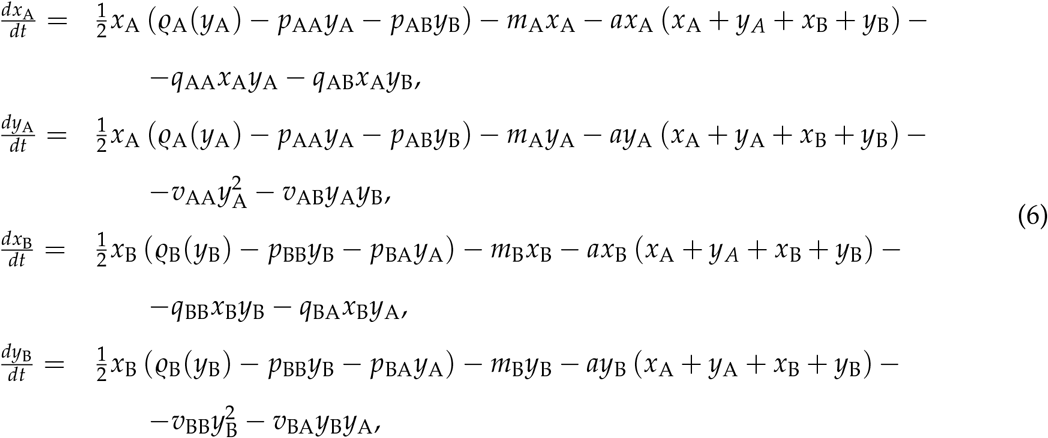

where the indices A and B distinguish between the species.

We are especially interested in coexistence of ecologically equivalent species. Therefore, we assume all parameters (ecological and sexual), except the *m*s, are the same for the two species. The coefficients of the intra- and interspecific sexual interactions (distinguished by subscripts w and b, as within and between) remain different. We allow *m*_A_ and *m*_B_ to differ, to test the robustness (structural stability) of coexistence against an advantage/disadvantage of one species relative to the other.

Note the non-triviality of the assumption, that the fertility functions of the two species are the same, despite their reproductive differences behind their genetic segregation.

For this case the ODEs read, as

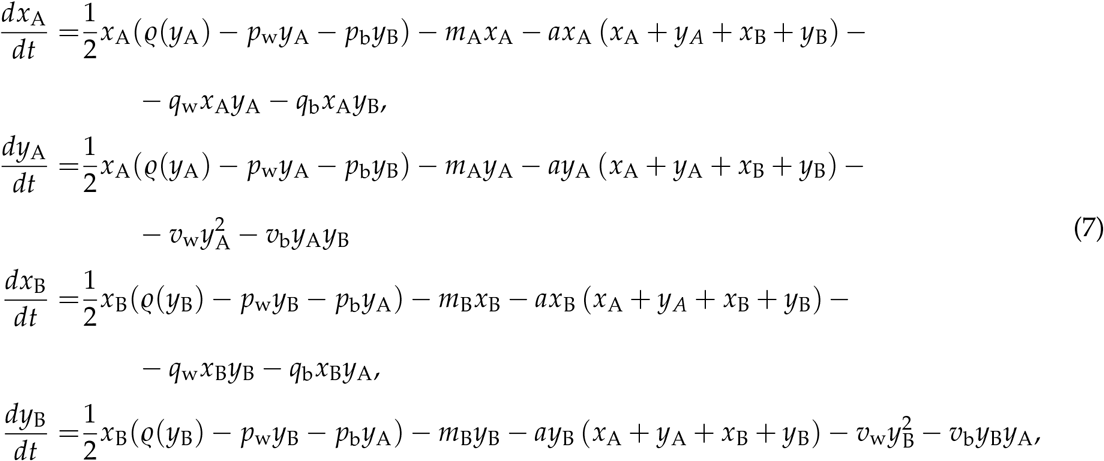

where *p*_w_ = *p*_*AA*_ = *p*_*BB*_ and *p*_b_ = *p*_*AB*_ = *p*_*BA*_; the same for the parameters *q* and *v*.

We will study the robustness of coexistence by varying the regulation-independent fitness difference (or, fitness difference, for short) Δ*m* = *m*_A_ − *m*_B_, while keeping 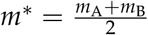 constant.

## Results

### Regulation of a single species

First we study, whether sexual interactions can regulate a single population. Figure 1 and 2 present phase portraits and time courses, respectively, for different cases and parameter combinations.

**Figure 1:**
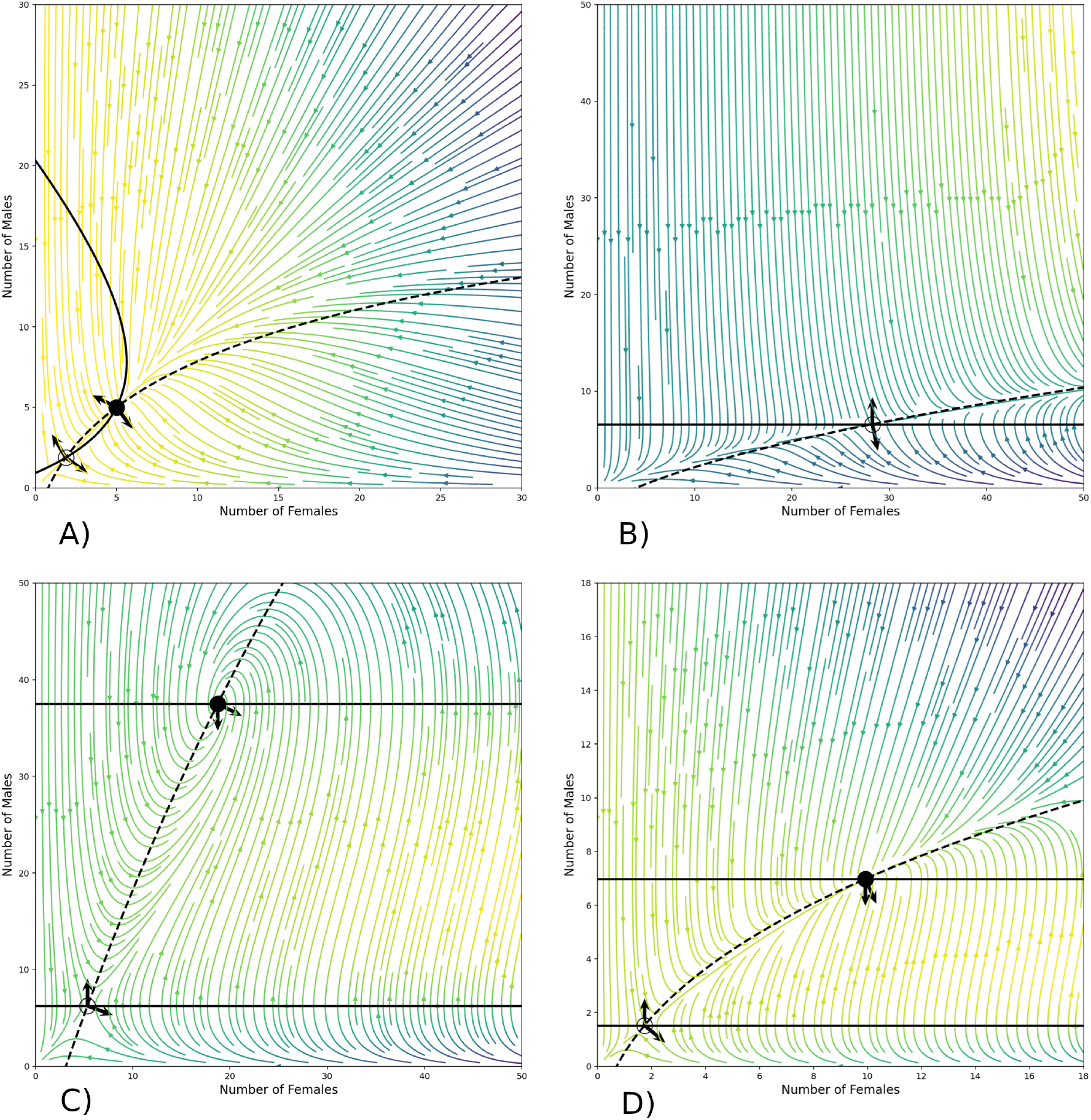
Phase portraits of the single species models. Solid and dashed black curves are female and male isoclines, respectively. Colored curves with arrows represent dynamics; lighter colors indicate higher speed. Stable and unstable fixed points appear as full and empty circles, respectively. Solid arrows show the gradients of the female and male dynamics at the fixed points. **A)** The two fixed points of the system with pure ecological regulation. (Parameters: *p* = *q* = *v* = 0, *m* = 0.2, 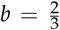, *T*_h_ = 0.02, 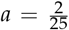) **B)** The single, unstable fixed point in absence of ecological regulation, with male competition only. (Parameters: *m* = 0.2, *v* = 0.1, *b* = 0.1, *T*_h_ = 0.9819, *a* = 0.) **C)** With female harm only: again two fixed points. (Parameters: *m* = 0.2, 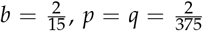, *T*_h_ = 0.8, *a* = *v* = 0.) **D)** The combined effect of the two sexual interactions. (Parameters: *m* = 0.2, 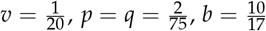, *T*_h_ = 0.8, *a* = 0.)

**Figure 2:**
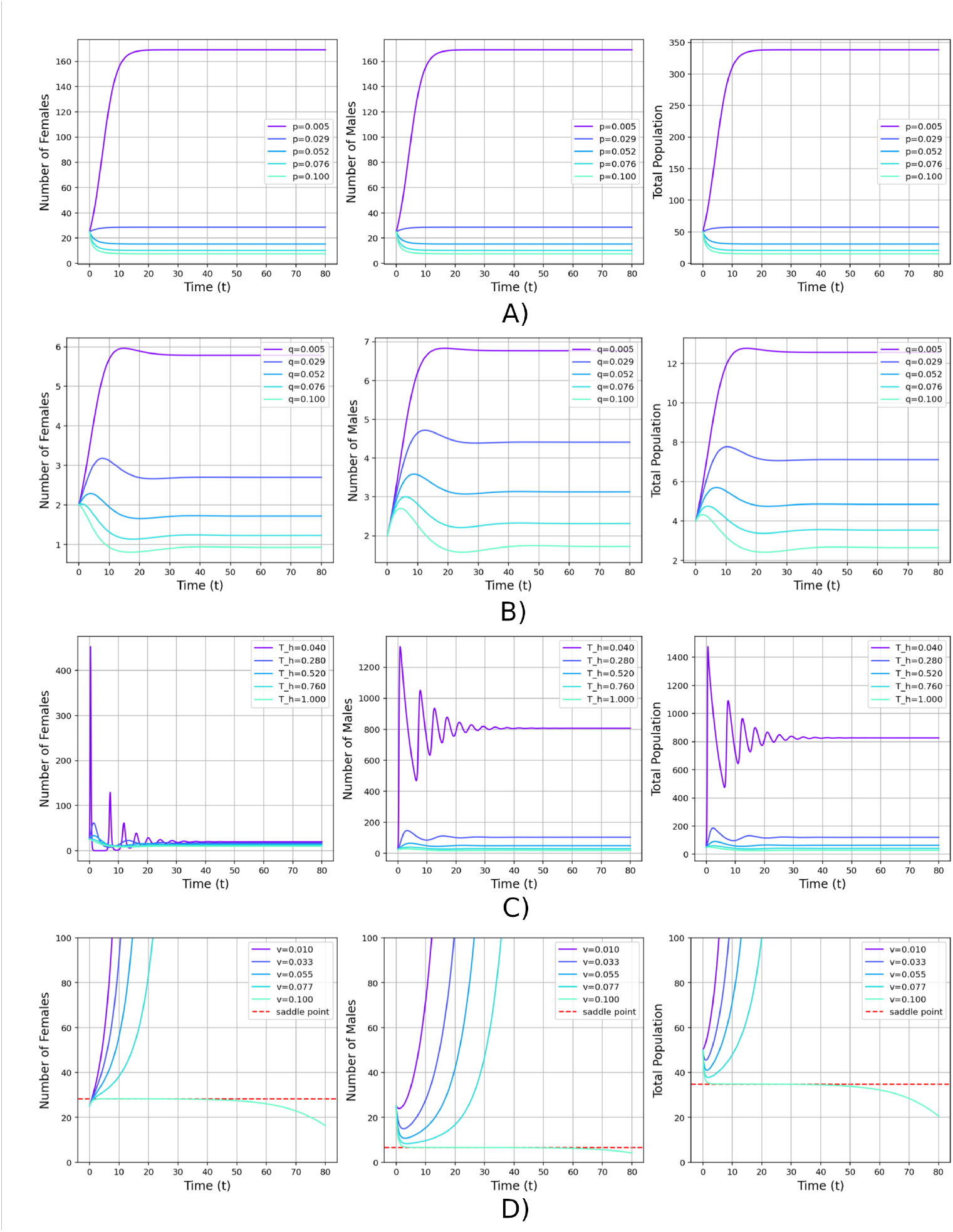
Time course of the single species model with different parameters in absence of ecological regulation. Both components of female harm stabilize population growth. **A)** Dependence on the male-induced decrease in female fertility. (Initial conditions: *x*_0_ = *y*_0_ = 25, parameters: *m* = 0.2, *q* = 0, *b* = 2, *T*_h_ = 0.8, *a* = 0; *p* varies.) **B)** Dependence on the male-induced increase in female mortality. (Parameters: *x*_0_ = *y*_0_ = 25, *m* = 0.2, *p* = 0, *b* = 2, *T*_h_ = 0.8, *a* = 0; *q* varies.) **(C)** Handling time length-dependence: transient oscillation with shorter handling time. (Parameters: *x*_0_ = *y*_0_ = 25, *m* = 0.2, *p* = *q* = 0.01, *b* = 2, *a* = 0; *T*_h_.) **(D)** Dependence on the strength of male competition: no stabilizing effect. (Parameters: *x*_0_ = *y*_0_ = 25, *m* = 0.2, *b* = 0.1, *T*_h_ = 0.9819, *a* = 0; *v*.)

Our reference case is a sexual population regulated solely by an ecological feedback. This system exhibits two fixed points as can be seen in Fig. 1A. Dependence of female fertility on the availability of males is a destabilizing factor at low abundances, resulting in an unstable fixed point at the Allee threshold. At higher abundances the fertility function saturates, but the population growth is limited by the total population size *x* + *y*, leading to another fixed point, which is stable.

In absence of ecological regulation, the presence of male competition alone (i.e., when *a* = *p* = *q* = 0, *v* > 0) is unable to constrain population growth. The system has only one, unstable fixed point in this case (Figs. 1B, 2D). While male competition has a negative effect on the abundance of the males, this affects female fertility and growth only through Allee effect. Therefore, male competition increases the Allee threshold for the total population, but does not result in a new, stable fixed point.

On the other hand, female harm, if present can regulate population growth (see Fig. 1C). We found that both components of female harm have the ability to stabilize even separately (see Fig. 2A and Fig. 2B).

Our results also showed that a decrease in handling time introduces a transitional oscillation into the system (see Fig. 2C and, for a detailed analysis, the Appendix 2). As the half-saturation level *k* appearing in female fertility (Eq. 5) is inversely proportional to *T*_h_, a longer handling time results in shifting the Allee threshold to lower population sizes. In case of a large *T*_*h*_, reproduction is not limited by searching for a mate, and the population growth is equivalent to an asexual one.

See Sections *Population dynamics of a single species* and *Transient oscillation* of the Appendix for analytic underpinning of these results.

### Coexistence of ecologically equivalent species

Based on the results on a single species, here we study the coexistence of two of them, as specified in the Model definition, especially Eq. (7).

In Figs. 3-5 the equilibrium states are plotted, as a function of Δ*m*, at different combinations of the sexual interactions. Because of the symmetry of the two species, the neutral coexistence point always exists for Δ*m* = 0.

**Figure 3:**
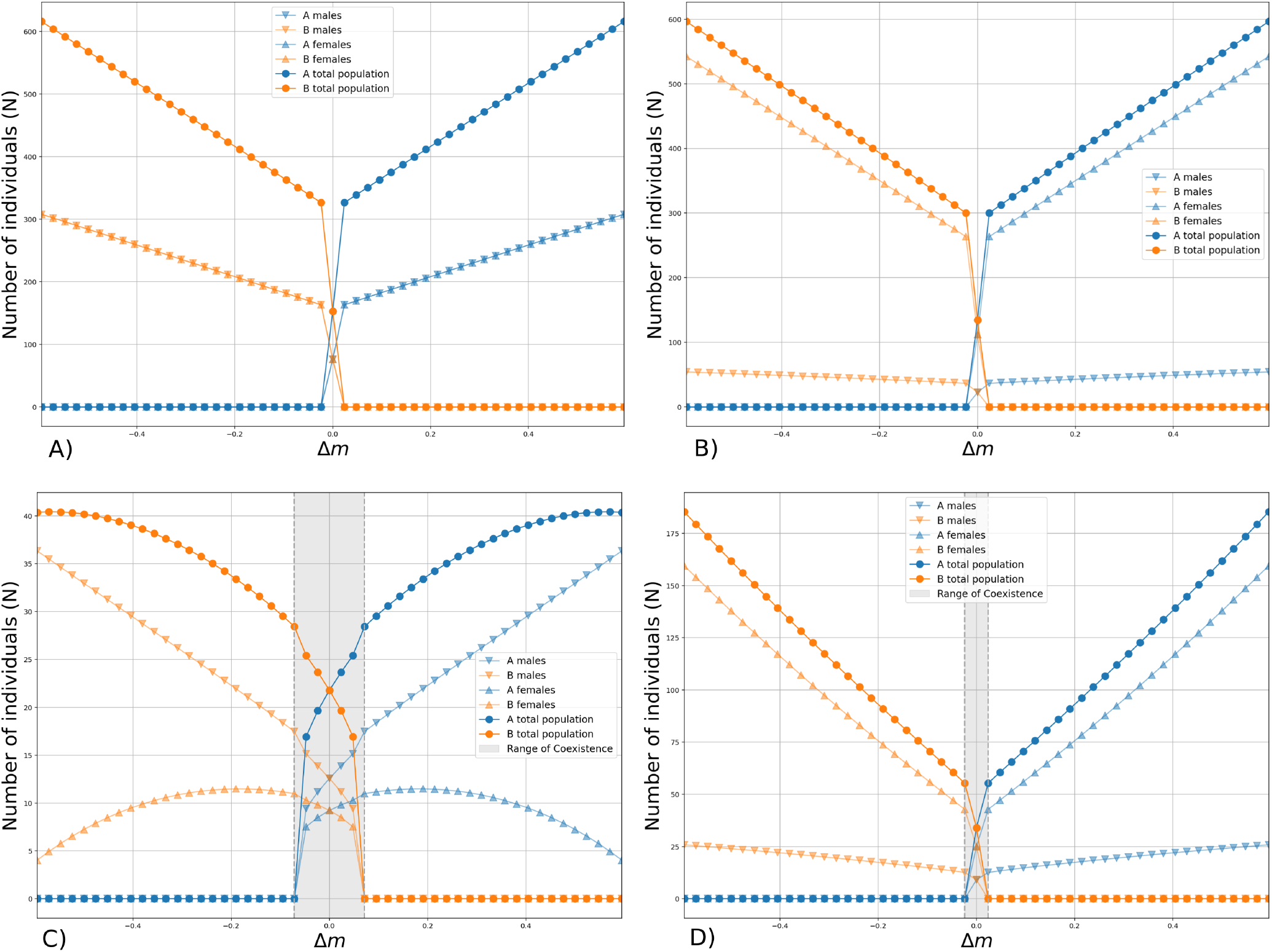
Robustness of coexistence with different kinds of regulations. Equilibrium population sizes of two ecologically equivalent species are plotted as a function of fitness difference characterized by Δ*m*. Range of coexistence – if present – is colored in gray. **A)** No sexual interactions: structurally unstable neutral coexistence. (The male and female abundances are equal. Parameters: 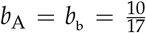, *T*_h,A_ = *T*_h,B_ = 0.8, *a* = 0.001 and *m*^∗^ = 0.3025. The same values are used for the rest of the two-species simulations.) **B)** Presence of male competition alone does not change the situation. (Additional parameters: *v*_AA_ = *v*_BB_ = 0.1, *v*_AB_ = *v*_BA_ = 0.) **C)** Intraspecific female harm alone: finite rage of coexistence. (Additional parameters: *p*_AA_ = *q*_AA_ = *p*_BB_ = *q*_BB_ = 0.01, *p*_AB_ = *q*_AB_ = *p*_BA_ = *q*_BA_ = 0.) **D)** Presence of both male competition and female harm: range of coexistence is shrunk relative to **C)**. (Parameters for the two kinds of sexual interactions are the same as in **B)** and **C)**.)

Without sexual interactions competitive exclusion is observed for any no-zero Δ*m* (Fig. 3A). Presence of male competition suppresses male abundances in both species, but does not eliminate competitive exclusion for any Δ*m* ≠ 0 (Fig. 3B).

In contrast, presence of intraspecific female harm stabilizes coexistence in a finite range of Δ*m*. Figure 3C demonstrates the effect with equal presence of the two harmful effects of female harm: diminished fertility and increased mortality. Figure 4 shows the stabilization by the two effects, separately. The coexistence range becomes narrower by the additional presence of male intraspecific male competition (Fig. 3D).

**Figure 4:**
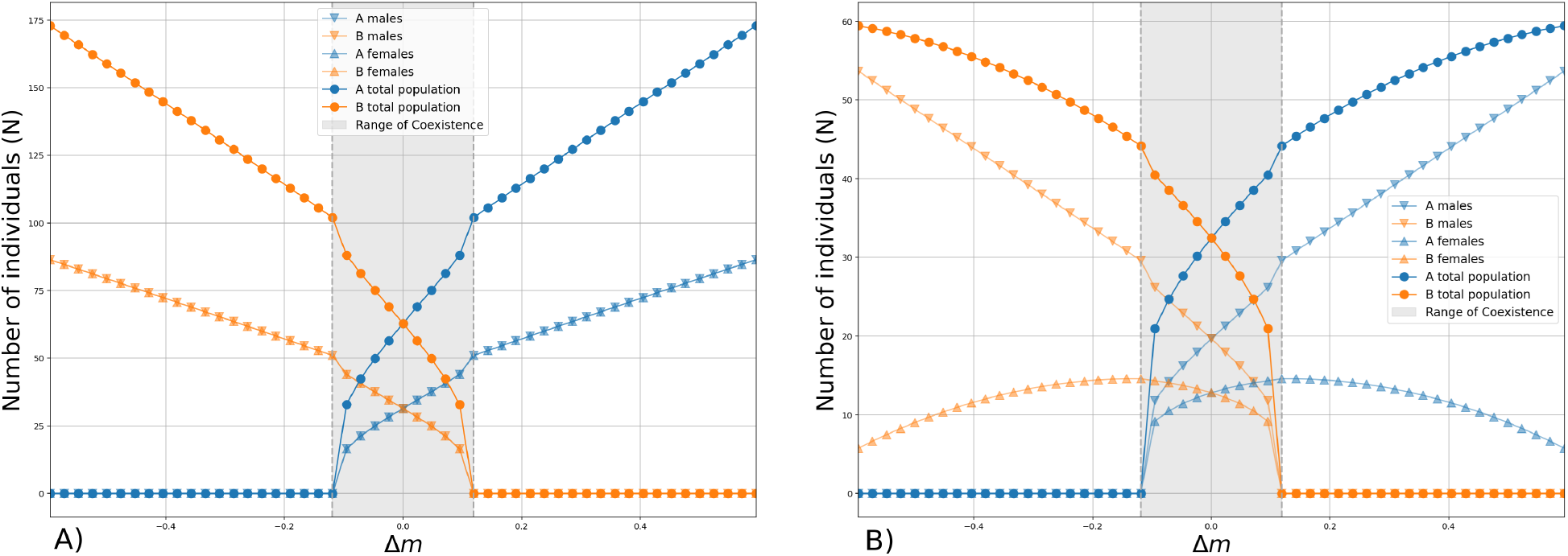
Effects of the two components of female harm, separately. **A)** Male abuse decreases female fertility. **B)** Male abuse increases female mortality. (Parameters, as in Fig. 3C, except that *q*_AA_ = *q*_BB_ = 0 in A and *p*_AA_ = *p*_BB_ = 0 in B.)

When both intra- and interspecific female harm is present, the the coexistence range is diminishing with strengthening interspecific harm (Fig. 5).

**Figure 5:**
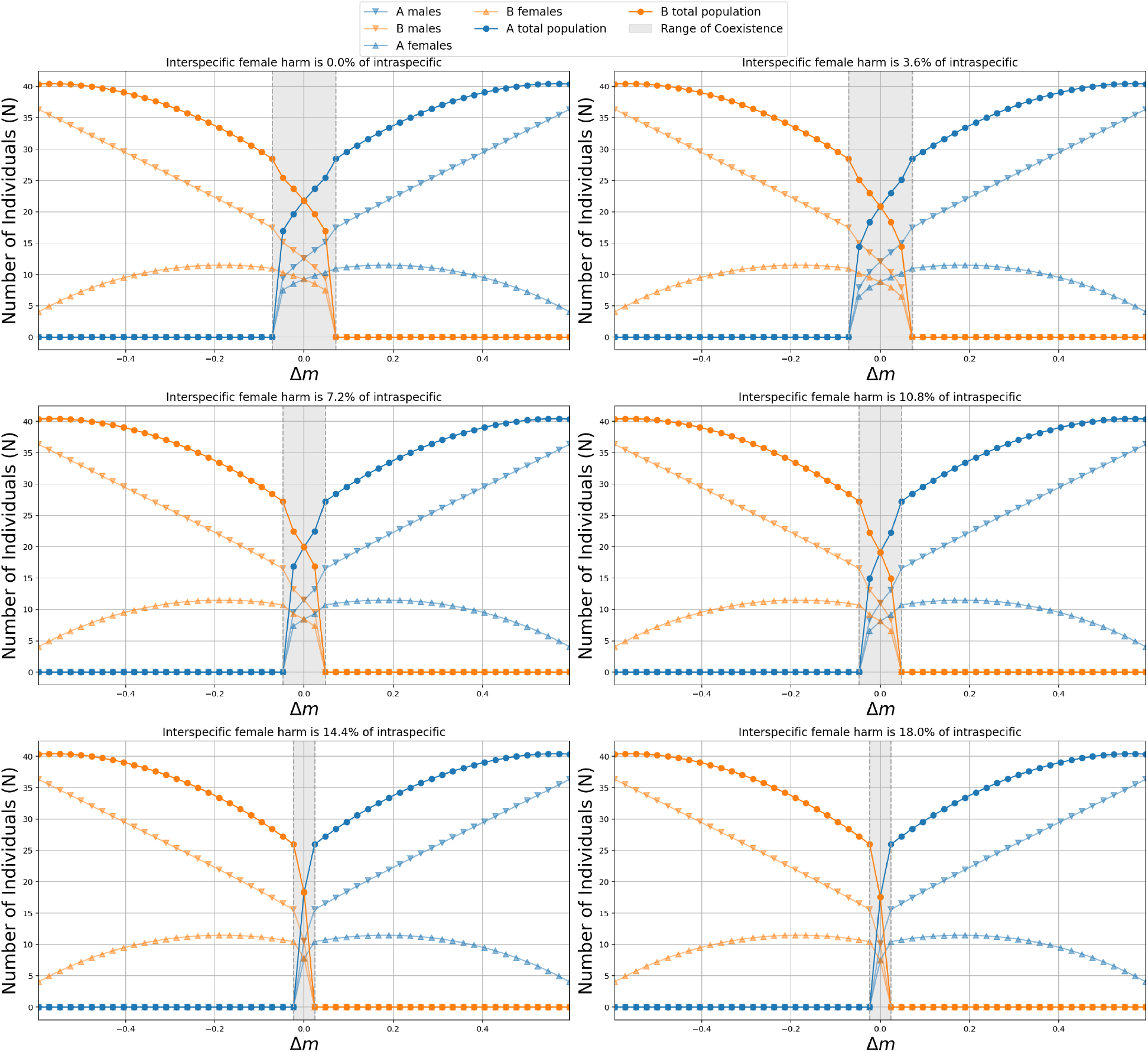
The effect of interspecific female harm on coexistence. As the strength of interspecific female harm increases, the range of coexistence shrinks. When the level of interspecific female harm reaches a critical value compared to that of intraspecific, the range of coexistence ceases to exist. (Parameters are the same as in Fig. 3C, but interspecific hams added to both components of female harm in percentage of the intraspecific harm.)

We conclude that intraspecific female harm is able to stabilize the otherwise neutral coexistence.

Section *Coexistence of ecologically identical species* of the Appendix contains the fixed point analysis, which determines the exact conditions of the dynamical and structural stability of coexistence.

## Discussion

We investigated two kinds of sexual interactions, female harm (a between-sexes interaction, Gómez-Llano et al. 2021, Figure 2) and male competition (a within-sex interaction), as proposed candidates for stabilizing coexistence of cryptic species pairs, in a homogeneous situation. We have learned that their effects are quite different, despite the parallelism of having a harmful intraspecific interaction in both cases. Female harm (diminished fertility, survival, or both) can limit population growth even without any ecological regulation. Relatedly, female harm is able to stabilize the coexistence of ecologically equivalent species, provided that intraspecific female harm is stronger than the interspecific one. In contrast, interspecific female harm destabilizes such coexistence. In this sense female harm behaves similarly to ecological regulation. Consequences of male competition is different. It is unable to limit population growth without other kinds of regulation. Accordingly, male competition does not stabilizes neutral coexistence even if it is purely intraspecific.

These conclusions underline the importance of mechanical modeling of specific interaction mechanisms – it was needed to recognize the essential difference between the consequences of the two kinds of intraspecific sexual interactions: female harm and male competition. Using the total population sizes, as dynamical variables, would mask this disparity. We were able to clarify also, that the two types of female harm, affecting fertility and survival of the females have the similar stabilizing effect.

### Female harm and male competition – empirical background

Elementary competition theory (MacArthur and Levins, 1967) states that stable coexistence requires weaker inter-than intraspecific competition. It is important to understand the two kinds of sexual interactions we studied from this perspective.

Female harm, when females suffer copulatory wounding or other forms of traumatic mating, is an embodiment of sexual conflict, driven by the difference in reproductive interests of sexes. Males can traumatize females due to either a mismatch between male and female reproductive morphology or an aggressive mating behavior. Both of these can decrease the fertility and/or lifespan of the females (Gómez-Llano et al. 2018; Zhang et al. 1998).

The incompatibility of the reproductive organs, as a form of prezygotic isolation, is particularly prevalent in the context of interspecific mating attempts (Reinhardt et al. 2015; Sota et al. 1998), while aggressive mating behavior can occur during both inter- and intraspecific ones. Some types of aggressive mating behaviors, like the slight poisoning of females by males during copulation (present e.g. in the case of fruit flies), are of stronger effect on within-species matings, as the appearance of these particular mechanisms requires the compatibility of reproductive organs.

The situation is even more complicated, however. When male-inflicted traumatization is frequent – as can be assumed in cases of intraspecific mating attempts of species prone to such behavior –, female subjects often exhibit a rapid adaptation process, leading to the development of various defensive mechanisms. These include the evolution of hard armors, as observed in bedbugs and spiders (Reinhardt et al. 2003; Rezác 2009). However, in cases of sporadic traumatization – which usually occurs when males of a species prone to aggressive mating behavior attempt to mate with females of a species not prone to that –, the females do not adapt to it, and consequently, when such instances arise, they incur significant costs. Empirical evidences support that in most cases, interspecific female harm is of more severe consequences due to this (Sota et al. 1998). The costs are often so high that they can act as a barrier to hybridization, and so, interspecific mating (Reinhardt et al. 2015).

That is, female harm can occur both within and between species; their relative strength depends on the biological details. Our theoretical study confirms the analogy between ecological competition and female harm: *if* the intraspecific harm is stronger than the intraspecific one, *then* a stabilizing feedback arises.

In contrast, male aggression towards conspecific males are always stronger than towards heterospecifics (Kohda 1995, 1998). The emerging negative frequency dependence was supposed to maintain diversity (Svensson et al. 2018). However, we found that male competition, in itself, is unable to stabilize coexistence under the homogeneity conditions of our study.

As we assumed spatial homogeneity, our result does not exclude the coexistence-maintaining mechanism based on territorial fights of males, as suggested by Mikami et al. (2004) and Kohda (1995, 1998). They showed that males may chase their conspecific competitors further away than heterospecifics. This effect leads to a patchy distribution of species, and allow them to coexist.

Such coexistence is a borderline case between the ecologically and the sexually stabilized coexistence. While it is based on a sexual interaction, it can be seen as competition for food *and* for territory. The within- and the between-species competition for the common food source is of the same strength. On the other hand, their competition for the territory is stronger within-than between-species. This possible consequence of male competition is not represented in our model.

### Coexistence theory background

The theoretical context of the problem of cryptic species is the meandering discourse about competitive exclusion (or, Gause’s) principle, often referred to, as niche theory. Should competitive exclusion be regarded, as the default relation between populations? Or, coexistence should be seen as the natural situation (Neil et al., 2009)? Even if it is true that “complete competitors cannot coexist” (Hardin, 1960), isn’t it a uselessly open-ended question to find an arbitrarily small, and any kind of, difference between them (Rosenzweig, 1995, p. 127)?

While MacArthur and Levins (1967) and Hutchinson (1957, 1959, 1978) provided any early formalization of niche theory, doubts about the generality of this picture was expressed early on, even by Hutchinson (1961) himself, and, more definitely, by Huston (1979, 1994). It was perceived by many, that Gause’s principle was incompatible by the huge species diversity we observe. Especially, because the principle was often identified with the statement that no more species can coexist, than the number of the resources. Ubiquity of fluctuations and disturbances were cited often to invalidate the principle.

Recent opinions seem to converge to the acceptance of competitive exclusion as the default behavior, even in presence of fluctuations. This renewed picture is often referred to as Modern Coexistence Theory (Barabás et al., 2018; Chesson, 1991; Chesson and Huntly, 1997; Chesson, 2000; Ellner et al., 2018; Godoy et al., 2014; Yamamichi et al., 2023). Chesson (2000) enlisted the major types of niche differentiations: the coexisting species may differ in resource use, in enemies, or in spatial/temporal arrangement. (This list goes back to Darwin, Pásztor and Meszéna, 2022.)

Contemporary Niche Theory (Chase and Leibold, 2003; Leibold, 1995) is a converging (Letten et al., 2017) approach based on Tilman’s resource competition model (Tilman, 1982). Beyond resource competition, they recognize the role of predation (apparent competition, Holt, 1977), and even the positive interactions (Koffel et al., 2021) in coexistence theory. The mechanistic coexistence modeling can be “recasted” often in terms of the original, MacArthur’s theory (McPeek, 2022, p. 364–381).

The deeper unification goes back to Levin (1970). He argued early on that instead of resource counting, one should count all the variables behaving mathematically, like a resource. Meszéna et al. (2006) demonstrated in a model-independent way that robust coexistence requires segregation in regulation: the species have to differ both in their impact on, and sensitivity towards the “regulating” (Case, 2001, p. 146, Krebs, 2014, p. 271) variables involved in the feedback loop of regulation. Increasing similarity in regulation decreases the parameter range allowing coexistence. This is the correct general formulation of both the limiting similarity expectation (Abrams, 1983; MacArthur and Levins, 1967) and Chesson’s equalizing vs. stabilizing effects (Chesson, 2000).

The connection between similarity and parameter-sensitivity answers Rosenzweig’s concern about the practical uselessness of the competitive exclusion principle. Regulatory differentiation must be biologically significant and probably detectable, otherwise coexistence would be at the mercy of very specific parameter combinations. As robustness against parameter perturbations is the hallmark of the non-neutral, stabilized coexistence, we investigated it, as a function of fitness difference Δ*m* in our numerical work.

The regulation approach is easier to connect to trait-based mechanistic modeling, goes beyond pairwise coexistence and unifies modeling of discrete regulating factors with continuous niche axes. See Barabás et al. (2012a, 2014); Pásztor et al. (2016); Szilágyi and Meszéna (2009a,b) for further developments and applications, including handlig heterogeneous and fluctuating environment.

When species differ in their regulation, frequency-dependence appears, as the regulating environment becomes dependent on the relative frequency of the species (Heino et al., 1998; Pásztor et al., 2016, p. 123). This frequency-dependence is always stabilizing (for two species: negative), if all interactions are competitive (negative-negative). In the general case, however, differentiation in regulation guaranties only the robust (structurally stable) existence of an all-positive non-neutral (hyperbolic) fixed point. As we saw in the current study also, the dynamical stability (which is a stronger property, than the structural one, see also *Coexistence of ecologically identical species* in Appendix) should be assessed separately (Koffel et al., 2021). A negative frequency-dependence stabilizes coexistence, while a positive one destabilizes it.

### Neutrality, the distinct limiting case

When the difference in regulation diminishes, the coexistence range shrinks. In the limiting case, the stabilizing feedback is lost, and the coexistence becomes structurally unstable. Such neutral coexistence disappears when one of the species acquires an arbitrarily small advantage/disadvantage relative to the other.

Based on the success of neutral theory in population genetics, Hubbel (2001) proposed to explain the observed huge species diversity by fitness equality, without any niche differences. However, exact fitness equality of distinctly different species is prohibitively unlikely (Purves et al., 2010). Still, neutral coexistence of physiologically equivalent species is possible. In this case, fitness equality is ensured by the equivalence itself, just like in the case of silent mutants in population genetics.

Neutral coexistence is always temporary, however, because the abundance ratio of the two species changes via random drift. By this reason, neutrality is not always considered, as coexistence (McPeek, 2022, p. 344-363). More operationally, one should compare the expected time to extinction of one of the species to the time-scale of interest: the estimated duration of coexistence at a given location, or the evolutionary divergence time between the species. The order of magnitude of expected lifetime of neutral coexistence is determined, as the product of the combined population size and the generation time (coalescence time, Fisher-Wright model, e.g. Pásztor et al., 2016, p. 243). This time-scale can be quite long for e.g. an insect population with large population size.

It was suggested, that sexually generated positive frequency dependence can also maintain regional coexistence of ecologically equivalent species by segregating them to different locations (Gómez-Llano et al., 2021). However, this mechanism only shifts the issue of coexistence to the regional/metapopulation level. If the species are neutral also in their competition for new patches, then the product of the number of local populations and the patch turnover time becomes the coalescence time. Therefore, it depends on the specific circumstances and numbers, whether positive frequency-dependence prolongs, or shortens coexistence of ecologically equivalent species.

The situation becomes more involved, when the localities are not isolated. In the model by M’Gonigle et al. (2012), instead of the patches, the vicinities of the local maxima of the carrying capacity function are occupied by the one, or the other species. While not tested in the model, we suggest that this coexistence would also become temporal, i.e. essentially neutral on the long time-scale, with time-varying carrying capacity function.

It was suggested also that slightly dissimilar species may also coexist in an almost-neutral way: a small niche difference may compensate for a small difference in fitness. This way we would have a niche-neutrality continuum (Adler et al, 2007), which was proposed, as the context of understanding cryptic species (Svensson et al., 2018).

However, the trait-based analysis (as opposed to work on the abstraction level of the Modern Coexistence Theory, e.g. Fig. 2e. in Yamamichi et al., 2023) leads to a different conclusion. The stabilizing effect of a small difference in the niche-related trait is quadratically small, too small to compensate a linear fitness difference (e.g. the figure in May, 1973, p. 158, or Fig. 4 in Szilágyi and Meszéna, 2009a; see also Song, et al., 2019). In the canonical example the strength of competition between two bird species, as a function of the beak size of one of them, has a maximum at equal beak sizes. Therefore, the stabilizing difference between within- and between-species competition is quadratic around the neutral situation.

The general mathematical reasoning goes back to the theory of adaptive dynamics (Geritz et al., 1998; Meszéna, 2005; Meszéna et al., 2005). In that context the statement is that similar species can coexist only at the “singular” points, where the fitness, as a function of the trait, has a maximum, or minimum – the latter case leads to evolutionary branching. This is the ultimate ecological reason of the discreteness of species: contrary to Roughgarden (1979, p. 534–536), a continuum of types cannot coexist (Barabás et al., 2012b, 2013a,b; Gyllenberg and Meszéna, 2005; Meszéna and Dieckmann, 2019).

### Coexistence stabilized by sexual regulation

As we demonstrated here, intraspecific female harm can also regulate a population and is a viable candidate for a coexistence-stabilizing mechanism. In contrast, we found that male competition in our context is unable to constrain an otherwise unregulated population, and accordingly, is unable to stabilize neutral coexistence. This may seem counterintuitive, as male competition is also an intraspecific negative interaction.

In some sense, it is just semantic that a feedback by sexual interactions are not considered, as ecological. After all, the presence of males is a factor of the outside word, i.e., of the environment, for a female individual. In this sense, one can consider the females, as the replicating agents and male abundances, as the regulating variables. How does an increased female abundance feeds back to female reproduction via this regulating variable? Males are produced by the females. Male abundance affects back to the females positively through Allee effect and negatively through female harm. The the first effect contributes positively, while the second one negatively to the overall feedback. That is, female harm, if strong enough to overcome the destabilizing Allee effect, is able to stabilize female abundance. Without female harm, however, the sexual feedback is necessarily positive. Male competition is unable to overcome this situation, because it is not in the female-male-female feedback loop. We would miss this distinction without the explicit consideration of the male and female dynamics and interactions.

If female harm, but not male competition, is able to regulate a single population, then it may be capable to stabilize the coexistence of two. The condition is that the intraspecific negative effect must be larger than the interspecific one – just like in the usual case of resource competition. This stabilization must overcome the destabilizing effect of Allee effect and an extra destabilization by the intrespecific harm (see Section *Coexistence of ecologically identical species* of the Appendix).

While we did not model here, several combinations of regulations are feasible. In principle, any number of species can coexist with sufficiently segregated female harm – even without any ecological regulation. (We assumed existence of a single, common ecological regulation.) It is also a possibility, that two species have a common ecological regulation and one of the species is regulated also, by female harm. In this case the species without female harm should have some other, non-regulating disadvantage. The critical issue is, that female harm can play the role of a distinct regulating mechanism.

### Criptic species – Why?

In principle, cryptic niche segregation, neutrality and regulation by sexual interactions are all possible explanations for coexistence of cryptic species. The question is, how can tell them appart in specific situations?

The simplest possibility is that cryptic species just coexist neutrally. While they necessarily differ in some sexual traits, this difference should not generate a difference in fitness. This can happen, when the fecundity functions of females are saturated, therefore female fertility does not depend sensitively on success of finding a mate.

A neutral species pair cannot arise from a niche-segregated pair by converging evolution, as suggested by McPeek (2019). The reason is that an arbitrarily small asymmetry/asynchrony between the converging evolutionary processes would prevent them to travel the full path to neutrality without experiencing competitive exclusion (Pásztor et al., 2020). Also, it is difficult to imagine their emergence via sexual selection, because it would assume non-saturated fertility function (but see van Doorn et al., 2004). Still, such species pair may arise from a single species via Mayr’s allopatric route (Mayr, 1942), when no further evolution happen after emergence of reproductive isolation.

If the time-scale of observed coexistence of our cryptic species is much longer than the relevant coalescence time, then we have to look for a stabilizing mechanism. The most customary assumption is an ecological niche segregation, which is “cryptic”, i.e., not obvious by some reason. For instance, if we do not observe niche differentiation in adult insects, then, maybe, they differ in their diet, or habitat of their larval state (ontological niche shift, see Szilágyi and Meszéna, 2009b about niche theory of structured populations). Such species pair can arise through ecological/adaptive speciation, which is increasingly accepted nowadays, as the main route for speciation (Dieckmann and Doebeli, 1999; Meszéna and Dieckmann, 2019; Nosil, 2012).

If we understand the trophic relationships, habitats and timing/seasonality of all life stages of the of interest sufficiently, and still do not find significant differentiation in regulation, then we can reject the possibility of ecological segregation with high confidence and look for a different stabilization mechanism.

We have learned that male-induced female harm is also a possibility for coexistence-stabilization. As we are not looking for an arbitrarily small effect, stabilization by female harm requires significant, and probably detectable, male-induced reduction of fertility, survival, or both. It is essential to establish, whether intraspecific female harm is significantly stronger, than the interspecific one.

A cryptic species pair, stabilized by intraspecific female harm may arise through the allopatric route, if female harm was strong in the parent species. They will coexist in a regulated way after recontact, if mating attempts of males are sufficiently intraspecific. Maybe, the new species can arise sympatrically also, as an escape from the female harm caused by the males of the the abundant parent species. Allee effect can be a barrier in the second scenario. These possibilities require further theoretical studies.

Several aspects of the topic have remained outside of our study framework. We assumed complete reproductive isolation between the species. Modeling speciation would require consideration of partial segregation and hybridization. Also, we assumed equal sex ratio at birth and equal density independent mortalities for the two sexes, despite some papers suggesting role of sex ratio in our subject matter (Zhang et al., 2004). Moreover, we assumed spatially homogeneous, sympatric, well-mixed populations. This assumption excluded us to study two possible effects: the spatial mosaic effect of male competition and the consequences of positive frequency-dependence.

Different pairs of cryptic species may have different coexistence mechanisms behind them. Empirical studies should have the last world. Species-specific empirical research can determine whether there is, in fact, no as yet unexplored segregation in the niches. Or, whether the equilibrium population size is, in reality, lower than would be justified by ecological regulation (which may indicate the presence of a different regulatory mechanism). Female harm itself is probably detectable. To exclude the possibility of neutral coexistence, time-scales and population fluctuations should be studied.

Good theoretical understanding of the possibilities may help to identify the specific mechanisms operating in specific systems on the basis of qualitative expert knowledge about the species in question. We hope that the current analysis contributes to the inquiry.

## Acknowledgments

We would like to express our gratitude to Tamás Székely for providing expert guidance in empirical results that enhanced our understanding of the actual biological processes underlying our analysis. We would also like to thank Justin S. Bois for the freely available Python function that acted as the core part of the phase plotter in our numerical analysis. This work was supported by the NKFI grant No. A152489.

## Statement of Authorship

A.F.F. and G.M. jointly conceptualized and developed the theoretical model and conducted the analytical analyses. A.F.F. performed all simulations and produced the figures. Both authors substantially revised the manuscript.

## Data and Code Availability

All code are available on GitHub (https://github.com/FFAdina/coex_sex).

## Appendix

### Population dynamics of a single species

Here we study dynamics of a single species, as defined by ODE (1). We rely on the general conditions for the fertility function *ϱ*(*x*), as specified in the model definition, but independent of any specific form, like Eq. (4). We will always assume that the parameters are such that the population is viable. We interested in the non-trivial fixed points only, so *x, y* > 0 is assumed.

We demonstrate the followings:

- An instable fixed point, related to Allee effect, always exists.
- Another, stable fixed point exists also in presence of ecological regulation, female harm, or both.
- Intraspecific female harm should be stronger than the interspecies one to stabilize coexistence. Allee sensitivity has a destabilizing effect.
- Consequences of the two kinds of female harm, affecting either the fertility, or the survival of the female, are similar.
- Male competition alone, without ecological regulation or female harm, is unable to constrain population growth.
- The stable fixed point is often reached thorough a transient oscillation. There is no limit cycle attractor in the model.

From Eq. (1), the equilibrium conditions are

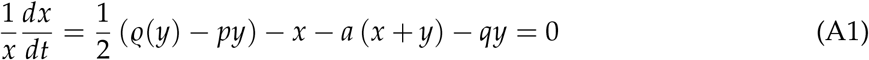

for the females (solid isoclines in Fig. 1, *x* ≠ 0 was used) and

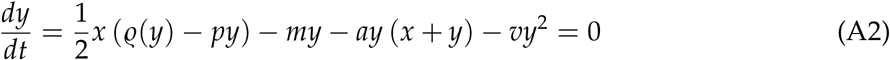

for the males (dashed isoclines). Observe that male competition does not affect female dynamics directly. This circumstance will be consequential.

Rearrangement of the differences of the female and male equilibrium conditions Eqs. (A1, A2) leads to the expression

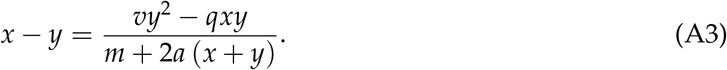

As we have equal sex ratio at birth, the equilibrium sex ratio is determined by the sex-dependent mortalities. Male competition suppresses the abundance of males, while male-induced female mortality suppresses the abundance of females. Sex ratio is unaffected by male-induced suppression of female fertility.

The male equilibrium condition (A2) determines equilibrium female abundance, as a function of the male one, uniquely:

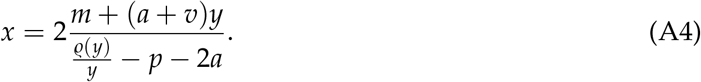

As the female equilibrium condition is more difficult to handle, it is worth considering two specific cases separately.

#### Equilibrium points in case of pure ecological regulation

In case of no sexual interactions (*p* = *q* = *v* = 0) Eq. (A3) implies *x* = *y* in equilibrium. Then substituting Eq. (A4) with *p* = *q* = *v* = 0 to Eq. (A1) leads to

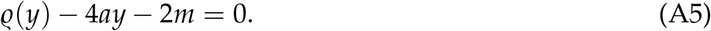

As *ϱ*(*y*) is an increasing, but saturating function, as specified in the Model definition, the l.h.s. of Eq. (A5) is an unimodal function for *a* > 0. Therefore we have at most two solutions for *y*; Eq. (A4) provides a single *x* value for each of them (see the two intersection points in Fig. 1A).

As we will see, the root with the lower *y* is the unstable Allee equilibrium, the other, stable one, comes from the ecological regulation.

In case of *a* = 0, *y* = *ϱ*^−1^(2*m*) is the only fixed point, corresponding to the Allee threshold. A population initiated above it grows unbounded.

#### Equilibrium points in the case of no ecological regulation

Here we assume no ecological regulation (*a* = 0), but allow sexual interactions. Then the female equilibrium reads, as

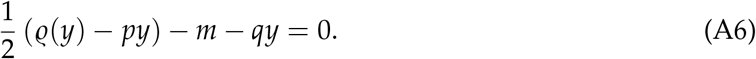

Compared to Eq. (A5), we can see that female harm works, as a substitute of the ecological regulation. The fertility and the mortality components (*p* and *q*) of female harm have identical consequences here. Again, we have at most two equilibrium points (Fig. 1CD); only one if *p* = *q* = 0.

As male competition does not enter to this equation, it cannot substitute female harm here. If neither ecological regulation, nor female harm is present, we have the *y* = *ϱ*^−1^(2*m*) Allee fixed point only, irrespective of male competition (Fig. 1B).

Also, male competition does not affect the equilibrium value(s) for male abundance *y*. The only effect of male competition is to increase the equilibrium female abundance *x*, according to Eq. (A3). Therefore, male competition has an effect of increasing total abundance of the population at the Allee threshold, as mentioned in the Results section.

Unfortunately, simultaneous presence of ecological regulation and sexual interactions does not allow a similarly simple analysis of the equilibrium points. However, the stability analysis below convince us the situation will not be more complicated, than the two nontrivial fixed point.

#### Stability analysis

Here we investigate the stability properties of the fixed points of the system (A1-A2) with full generality, i.e. allowing presence of ecological regulation and sexual interactions.

The Jacobian matrix elements at the fixed point of interest reads, as

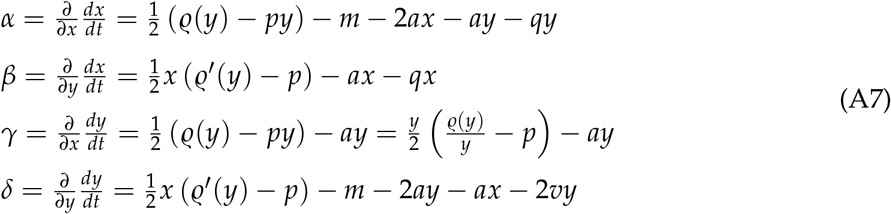

The eigenvalues of the Jacobian are

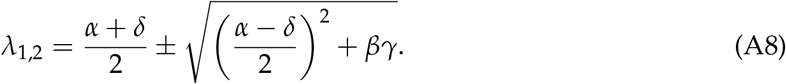

When the second term is real, then the sign of *βγ* decides whether the two (real) eigenvalue have the same, or the opposite, signs. In case of an imaginary second term the sign of the first term determines the stability/instability of the focus point.

The matrix elements are affected by the equilibrium conditions. The female equilibrium (A1) implies, that

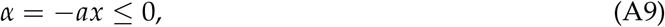

and

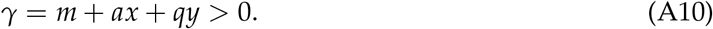

The male equilibrium (A2), combined with inequilibrium (2) leads to the upper bound

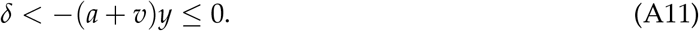

As *α* + *δ <* 0 and *γ* > 0, the nature of the stable point is determined solely by *β* (Figure A1):

- 0 *< β*: Two real eigenvalues with different signs, i.e., a saddle point;
- 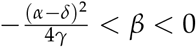: Two negative real eigenvalues, i.e., stable node;
- 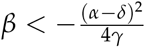: Complex eigenvalues with negative real parts, i.e., stable focus.

Therefore, the fixed point is stable, when *β <* 0, and unstable, when *β* > 0. That is, the stability condition can be written, as

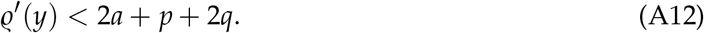

The Allee sensitivity term *ϱ*′(*y*) describes destabilization by Allee effect. The terms of the r.h.s. represent the stabilizing effects: the ecological regulation and the two types of female harm. Observe that male competition does not enter into the stability condition.

**Figure A1:**
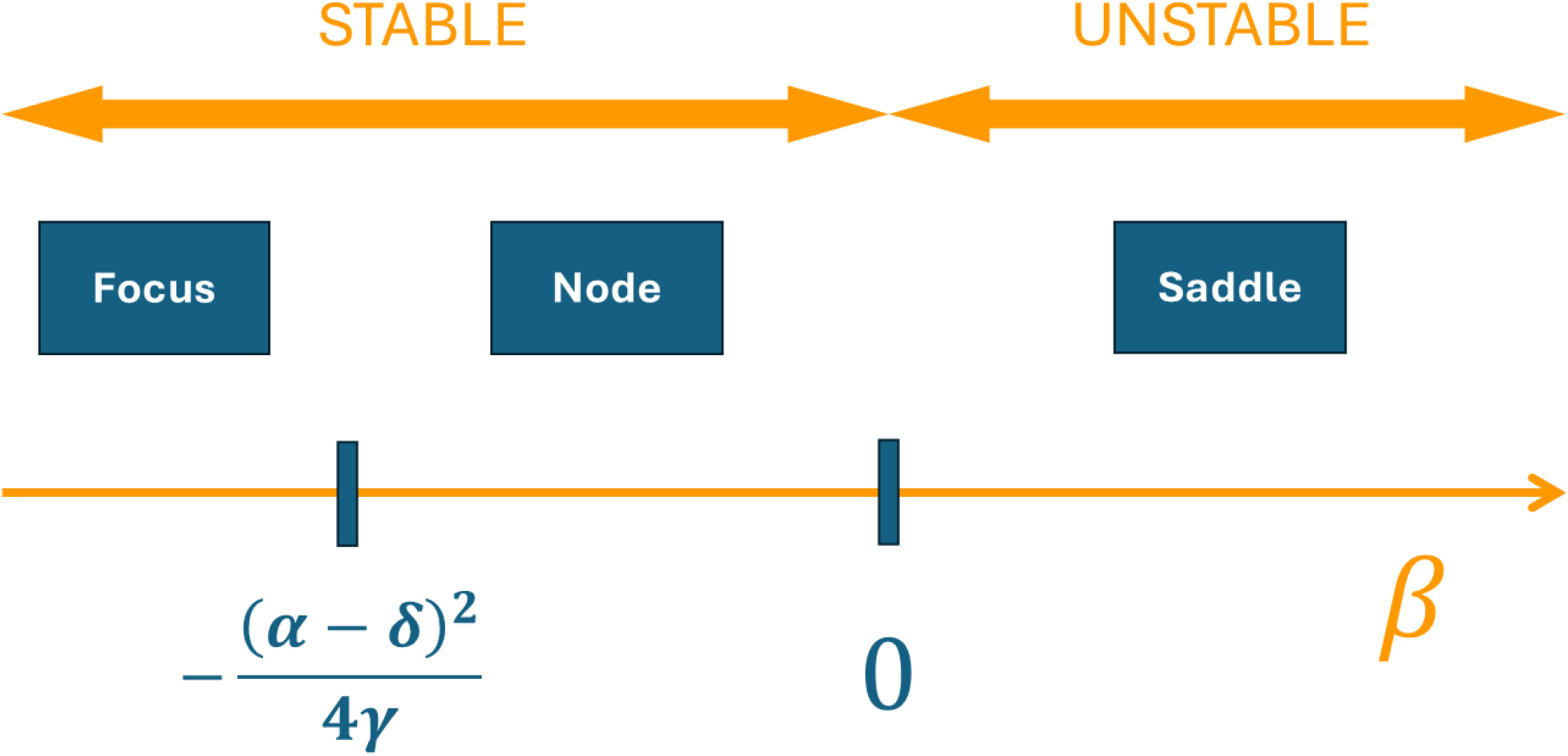
Classification of the fixed points. The properties of the fixed point is determined by the eigenvalues (A8) with assuming that *γ* > 0 and *α* + *δ <* 0. These assumptions hold for the stability analysis of a single population in Sectinon *Population dynamics of a single species* of the Appendix. The figure remains applicable for the two species analysis in Section *Coexistence of ecologically identical species* of the Appendix, with the parameter change 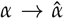, etc., when 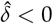, which is the more probable case.

As *ϱ*′(*y*) is a decreasing function, Eq. (A12) means that a fixed point is unstable below a critical value of *y* and stable above it. (We assume that *ϱ*′(0) is large enough, otherwise the population could not exist.) In presence of neither ecological regulation, nor female harm, i.e., when *a* = *p* = *q* = 0, *β* is always positive and there is no stable fixed point. Male competition can have some stabilizing effect through making *δ* more negative, but it is unable to constrain population growth in itself.

This result proves that no more than two non-trivial fixed point, an unstable, and a stable, can exist even in the joint presence of both ecological regulation and sexual interactions. The critical value of *y* divides the male isocline (A4) into an unstable and a stable arc. However, the successive fixed points should be alternately stable and unstable along the isocline – which exclude having more than two fixed points.

If the abundance at the stable equilibrium point is much higher, than the Allee threshold, then the Allee effect becomes negligible and *ϱ*′(*y*) ≈ 0 makes the stability condition (A12) very simple: intraspecific female harm must be stronger, than the interspecific one.

No parameter combination leads to an unstable focus point, which would be indicative for existence of a limit cycle. A more complicated attractor is not possible in 2D.

### Transient oscillation

As we have learned in Section *Population dynamics of a single species* of this Appendix, the case 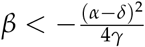 leads to a stable focus, i.e., the equilibrium point is reached via a transient oscillation, as observed in Figure 2. The figure shows that the amplitude of the oscillation decays faster for longer handling time *T*_h_. Here we reproduce and support this observation analytically. We will need the explicit formula (4) for the fertility function.

The angular frequency *ω* of the oscillation and the rate *A* of the exponential change of its amplitude correspond to the imaginary and the real part of the eigenvalue of the Jacobian matrix:

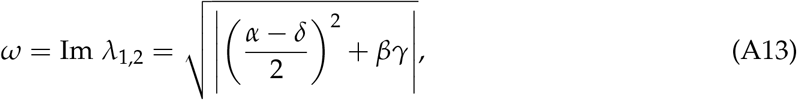

and

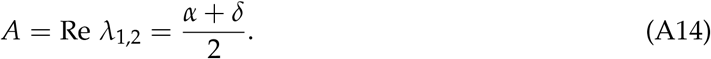

We are interested in the *T*_h_-dependent amplitude-change,

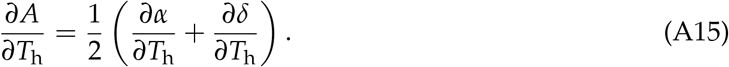

The elements of the Jacobian matrix depend on *T*_h_ merely through the fertility function and its derivative, so it is sufficient to look at the *T*_h_-dependence of these terms only, which will read:

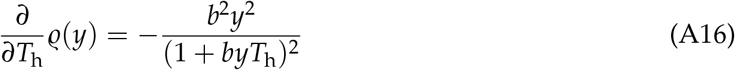

and

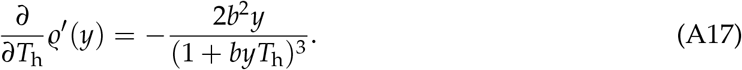

This way we can write the partial derivatives of the Jacobian matrix elements with respect to the handling time as:

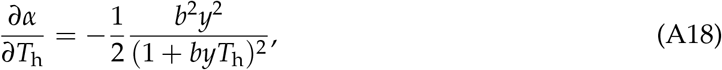

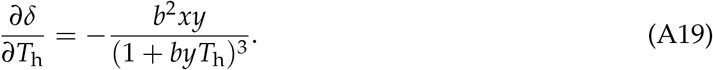

Substituting these into Eq. A15, we get the *T*_h_-dependence of the rate of the exponential change of the oscillation’s amplitude:

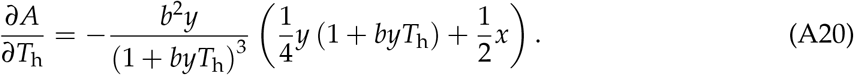

As 1 + *byT*_h_ > 0, the r.h.s. of Eq. (A20) is negative, so we can conclude that the handling time plays a damping role in the oscillation, i.e. an increase in the value of *T*_h_ will result in a more rapid decay of the oscillation. This result is understandable, because in case of large *T*_h_ reproduction is not limited by searching for mate. Population growth is equivalent to an asexual growth.

### Coexistence of ecologically identical species

Here we study the coexistence of two ecologically identical species with common ecological regulation, as specified by Eq. (7). Again, we assume a generic fertility function *ϱ*(*y*). All the parameters of the two species are the same, except the density independent mortality *m*. The potential difference between the *m*s will allow us to investigate robustness of coexistence against an advantage/disadvantage of one species relative to the other.

We will demonstrate, that dominance of intraspecific female harm over the interspecific one should overcome destabilization by Allee effect and an extra effect of very strong interspecies female harm.

#### Perturbation of the neutral coexistence

To make the symmetry and the differences explicit, we introduce the sum and difference variables:

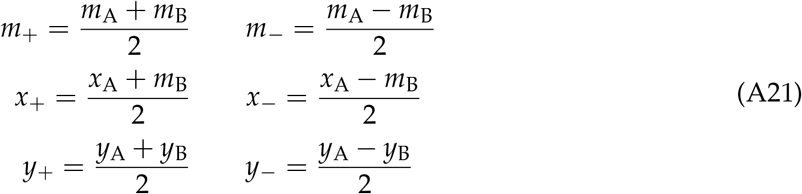

leading to *m*_A_ = *m*_+_ + *m*_−_, *m*_B_ = *m*_+_ − *m*_−_ and the analogues for the abundance variables. (Note that Δ*m* = 2*m*_−_ was used as parameter in the Figures.) Positive abundances imply

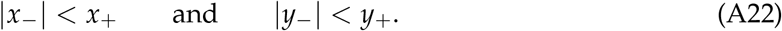

With this reparametrization of the dynamics (7) reads, as

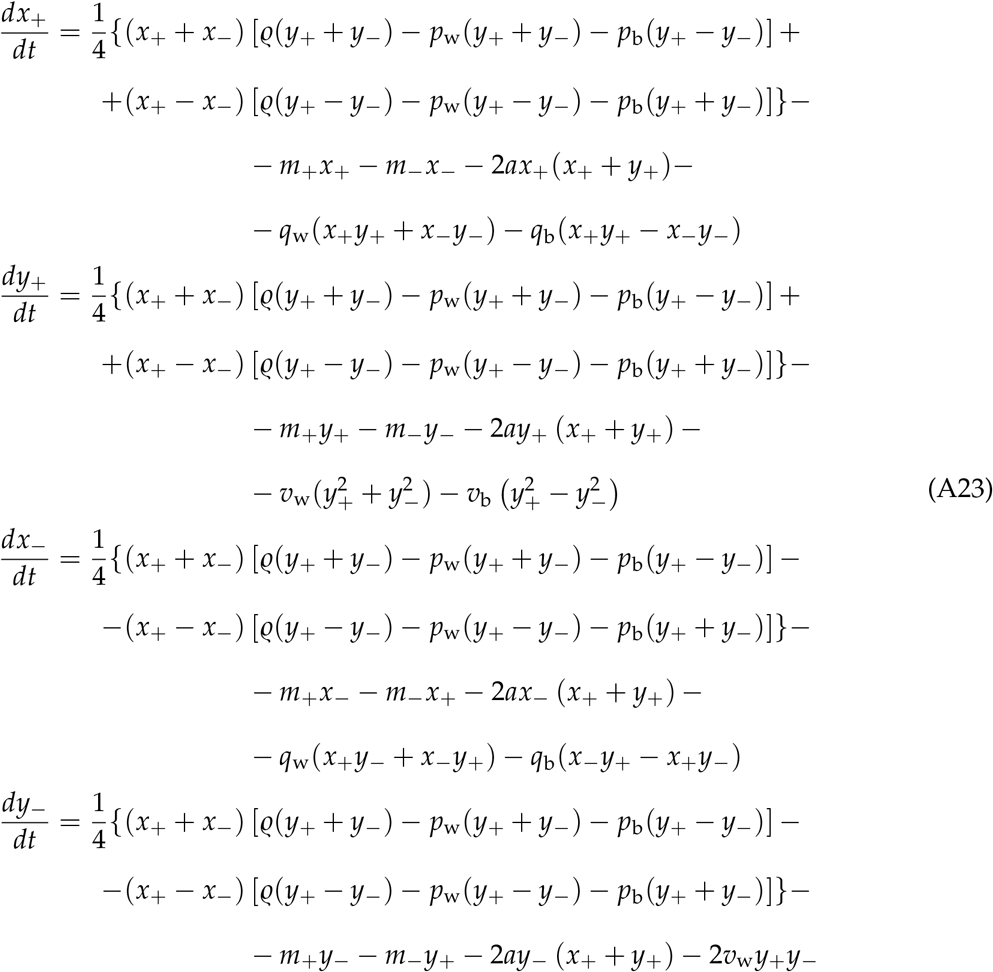

Assume first, that *m*_−_ = 0, i.e., that the two species are ecologically equivalent. Then, obviously, there is an equilibrium point with equal densities. Substituting *x*_−_ = *y*_−_ = 0 to the r.h.s. of the equations leads to

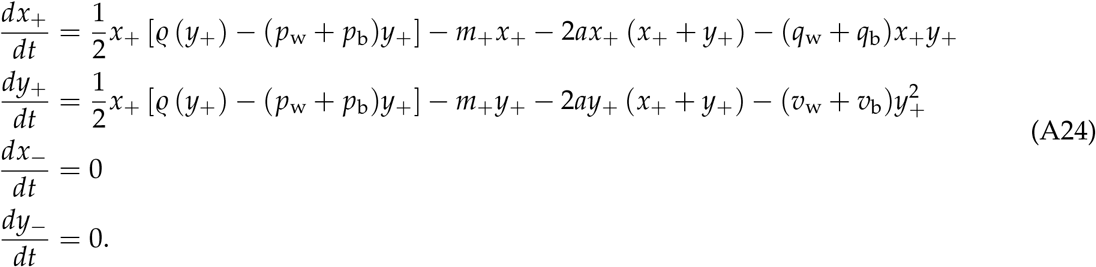

If both the female and male densities are equal for the two species, then it remains so forever.

The dynamics of the total abundances is like Eq. (1) for a single population. The equilibrium conditions for the total abundances read, as

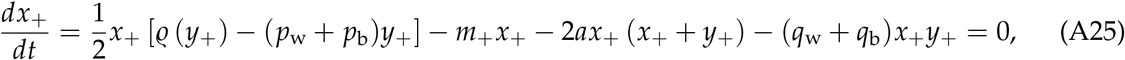

and

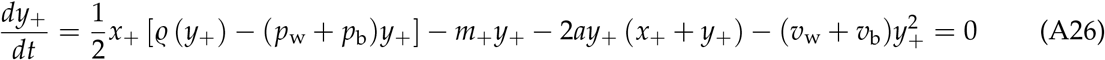

for the females, and the males, respectively.

To study the perturbation of this symmetric (neutral) equilibrium we linearize the dynamics (A23) in the small quantities *m*_−_, *x*_−_ and *y*_−_:

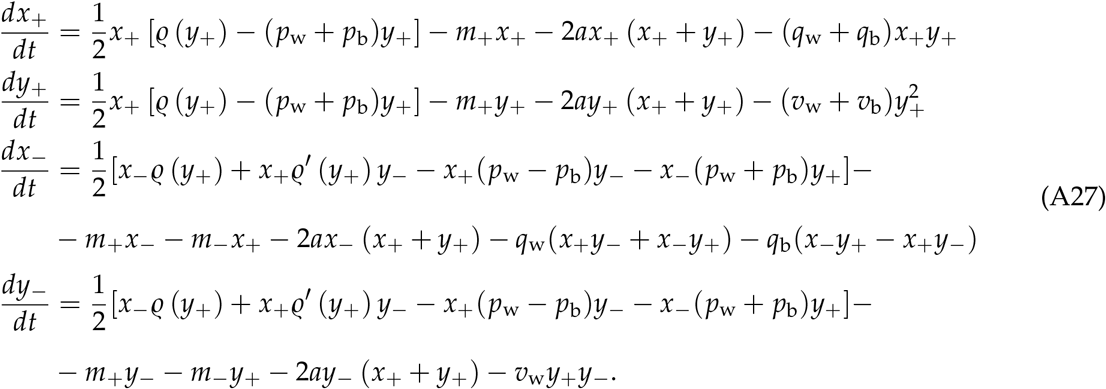

Observe, that in first order, the dynamics of the total densities are unaffected by the small difference quantities. Therefore, we can study the perturbation of the symmetric equilibrium in terms of the difference quantities, assuming that the total densities are in the equilibrium defined by the equations

The linearized difference dynamics can be written in the matrix form

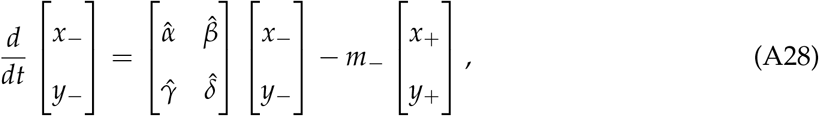

where

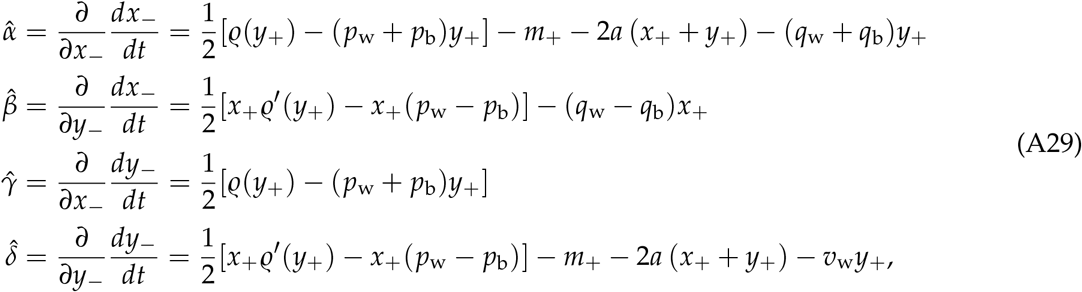

the derivatives are taken at *m*_−_ = *x*_−_ = *y*_−_ = 0.

Again, analogously to Eq. (A8), the eigenvalues of the Jacobian reads, as

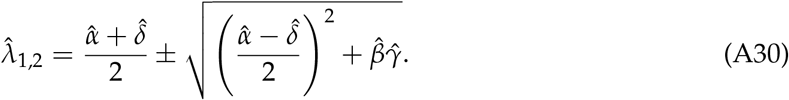

The equilibrium solution reads, as

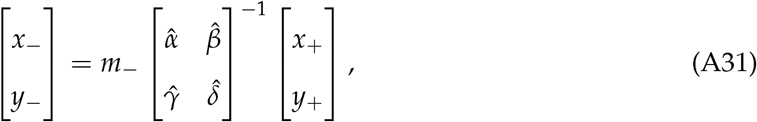

provided, that the matrix is invertible, i.e., none of the eigenvalues are zero. If this condition is fulfilled, the solution is structurally stable. Dynamical stability, which is a stronger property, than structural stability, requires negativity of both eigenvalues.

Note that Eq. (A31), being a linearization, is valid only, if |*m*_−_| is sufficiently small.

With increasing |*m*_−_| coexistence (gray regions in Figs.) disappears, when positivity conditions (A22) are falsified. While the extinction boundary cannot be calculated exactly from the linear approximation (A31), it is informative for qualitative understanding. When the stabilization becomes weaker, and one of the eigenvalues of the Jacobian matrix approaches zero, then the system becomes very sensitive to *m*_−_, and the extinction threshold will be reached at a low value of |*m*_−_|.

As an example, Fig. 5 demonstrates a shrinking range of coexistence, as increasing interspecific female harm weakens stabilization. Fig. 3D shows that presence of male competition shrinks the coexistence range stabilized by female harm. This happens because male competition suppress male concentration *y*_+_ and, consequently, shrinks the allowed range for |*y*_−_|.

#### Stability condition

To understand stability of the coexistence, we should calculate the elements of the Jacobian matrix at the fixed point.

The female equilibrium condition (A25) leads to

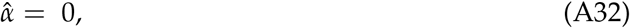

while either of the equilibrium conditions implies

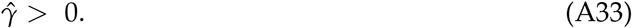

Finally, it follows from the male equilibrium (A26), that

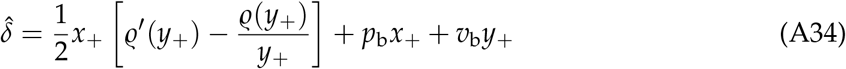

where the bracketed term is negative by inequality (2).

Because of the relations (A32-A33), negative definiteness has two conditions: (A) 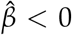 and (B) 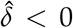. Positive 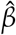 would result in two real eigenvalue with different sign, while positive 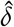 would lead a positive sum of the eigenvalues.

Condition (B) has a good chance to be fulfilled. The bracketed term in Eq. (A34) is negative by inequlibrium (2). Therefore 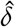 can be positive only, if the interspecies sexual effects are sufficiently large. Also, if *y*_+_ is sufficiently large and fertilitiy is saturated, therefore Allee sensitivity diminishes (*ϱ*′(*y*_+_) ≈ 0), then the rest of the term together is negative, as one can see from the male equlibrium (A26). That is, both the term *ϱ*′(*y*_+_) and the interspecific terms should be sufficiently large to make 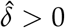.

If condition (B) is satisfied, i.e., 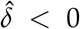, then the situation becomes similar to the one in Appendix 1, as pictured in Fig. A1. Then the stability is determined by condition (A), which can be written, as

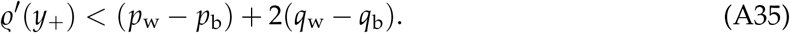

Note the similarity of condition (A35) to the stability condition (A12) for single species. In both cases, Allee sensitivity is the destabilizing force. Stabilization of coexistence is based on *differential* regulation of the species. Therefore the female harm terms of Eq. (A12) is replaced by the intra-inter differences in Eq. (A35). The ecological regulation does not contribute to the coexistence stabilization, because we assumed a common regulation.

As in the single species case, the two kinds of female harm, decreased fertility and increased mortality, have the same kind of stabilizing effect; male competition has no such effect.

If the equilibrium abundance is much higher than the Allee threshold, then the Allee destabilization becomes negligible. Then condition (B) is satisfied and the stability is governed solely by the relative strength of within- and between-species female harm.

If the Allee sensitivity is stronger at lower abundance, then both conditions of stability are relevant. Then, strong interspecific sexual interactions can destabilize coexistence also, independently of the strenth of the intraspecific ones. As an example, Fig. 5 demonstrates the shrinking of the coexistence range, when increasing interspecific female harm weakens stabilization.

